# IGPR-1 is phosphorylated on the immunoreceptor tyrosine-based motif, stimulates AKT pathway and supports melanoma growth

**DOI:** 10.1101/2024.08.19.608690

**Authors:** Nader Rahimi, Sreesaisowmya Potluri, Vipul Chitalia

## Abstract

Melanoma is a lethal form of skin cancer that impacts one out of every five Americans and ranks as the fifth most prevalent cancer among men and women in the United States. Immunoglobulin (Ig) and Proline-rich Receptor-1 (IGPR-1, also called TMIGD2/CD28H) is closely related to immune checkpoint, CD28/PDL1 family receptors. It controls important cellular processes including, immune cell regulation, cell-cell adhesion, mechanosensing, autophagy, and angiogenesis, and its activity is associated with multiple human malignancies. However, the role and signaling mechanism of IGPR-1 in melanoma remains largely undefined. Here, we report that IGPR-1 is mutated or upregulated in nearly 13% of melanoma and its pro-tumor signaling in melanoma cells is mediated by phosphorylation of immunoreceptor tyrosine-based activation motif (ITAM) tyrosine (Y222). IGPR-1 is phosphorylated at Y222 in human melanoma and cell culture. Phosphorylation of Y222 is context-dependent and is catalyzed by EGFR and Src kinase. Inhibition of EGFR by pharmacological and shRNA strategies inhibited phosphorylation of Y222, whereas stimulation with EGF promoted phosphorylation of Y222 in vivo and recombinant active EGFR catalyzed its phosphorylation in an *in vitro* kinase assay. *In vivo* co-immunoprecipitation and *in vitro* GST-pull-down assays demonstrated that phospho-Y222 facilitates the binding of IGPR-1 with the SH2 domain-containing proteins, SHC1 and SHP2. IGPR-1 stimulates multiple key signal transduction pathways relevant to tumorigenesis, including AKT, mTOR, and MAPK. Mutation of Y222 blocked IGPR-1-mediated activation of AKT and MAPK leading to inhibition of 3D-spheroid tumor growth. By investigating the immunoreceptor tyrosine-based motif signaling of IGPR-1, this study uncovers new findings that could have significant diagnostic and therapeutic implications for melanoma.

## Introduction

Immunoglobulin (Ig) and Proline-rich Receptor-1 (IGPR-1, also called TMIGD2 and CD28H) is a new member of the CD28 family receptors, which include CD28, cytotoxic T-lymphocyte–associated antigen 4 (CTLA-4), inducible co-stimulator (ICOS), program death-1 (PD-1), and B- and T-lymphocyte attenuator (BTLA)^1–2^ and is expressed in various human cell types including, immune cells, epithelial cells, endothelial cells, and skin melanocytes^2–3^. IGPR-1 regulates key angiogenic responses such as capillary tube formation and barrier function^2, 4–5^. IGPR-1 is therapeutically relevant receptor since it is upregulated in several human cancers including, colon cancer, gastric cancer, and acute myeloid leukemia (AML)^6–8^. It also has a prognostic significance because its overexpression is linked to poor survival ^6–7, 9^. At the cellular level, IGPR-1 senses and reacts to cellular stress by acting as a mechanosensing receptor^4–5, 9–10^, promotes cell survival through cell-cell adhesion, multicellular aggregation and autophagy^4–5, 10^.

The extracellular domain of IGPR-1 contains a single immunoglobulin (Ig) domain followed by a single transmembrane domain and a proline-rich (PXXP) intracellular domain. While the extracellular domain of IGPR-1 mediates homophilic dimerization and interaction with HHLA2, its PXXP cytoplasmic domain is subject to various posttranslational modifications (PTMs) including, phosphorylation and ubiquitination^5, 11–12^ and mediate protein-protein interaction by recruiting the Src-homology domain 3 (SH3) and WW domain-containing proteins such as SPIN90 and NEDD4 to IGPR-1^2, 11^. In response to cell-cell adhesion and cellular stress (*e.g*., shear stress, nutrient deprivation, and DNA damage causing chemotherapeutic agent doxorubicin), IGPR-1 is strongly phosphorylated at Ser220, and its phosphorylation is catalyzed by AKT/PKB and IKKβ^9–10^. The cytoplasmic tail of IGPR-1 also contains two putative immunoreceptor tyrosine-based activation motif (ITAM) tyrosine phosphorylation sites (VLY_197_RPR and SIY_222_STS). However, the role of these tyrosine phosphorylation sites in the IGPR-1 function remains undefined. In this study, we demonstrated that IGPR-1 is phosphorylated on Y_222_ in a context-dependent manner and identified EGFR and Src as the kinases responsible for catalyzing the phosphorylation of Y_222_. Phosphorylation of Y_222_ recruits SHC1 scaffold protein and SHP2 tyrosine phosphatase to IGPR-1 via the Src-homology domain 2 (SH2), stimulates activation of AKT and MAPK pathways and 3D-spheriod tumor growth.

## Materials and Methods

### Plasmids, antibodies, shRNAs and recombinant proteins

EGFR/ErbB1 and ErbB2 constructs were cloned into pLNCX retroviral vector were originally provided by Professor David F Stern, Department of Pathology, Yale University School of Medicine, New Haven, Connecticut. Recombinant human active EGFR protein (cat# ab155639) and recombinant active Src protein (ab# 60884) were purchased from Abcam, which were produced in Baculovirus system. GST-C-terminus IGPR-1 was previously described^11^. IGPR-1 constructs including, full length IGPR-1, N-terminus deleted and Y222 mutant IGPR-1 were cloned into a retroviral vector. GST-SH2 SHC1 (#46486)^13^ and GST-SH2 (encompassing both the N- and C-terminus) SHP2 (#67575)^14^ were purchased from Addgene. Phospho-Y222 antibody was developed using peptide CDQRGQSI-pY_222_-STSFPQP. Human lentiviral EGFR shRNA (cat# sc-29301), which is consist of pools of three to five target-specific 19-25 nucleotide sequences in length, was purchased from Santa Cruz biotechnology.

### Retrovirus production and transduction

Retrovirus particles were produced in 293-GPG cells, and viral supernatants were collected as previously described^15^ and used to infect melanoma cells or HEK-293 cells.

### Cell transfection

HEK-293 cells stably expressing IGPR-1 were transfected with EGFR or other constructs via PEI (polyethylenimine) as described^11^. After 48 h transfection, cells were stimulated with EGF, lysed, and subjected to immunoprecipitation or western blotting as described in the figure legends.

### Cell culture

Pathogen-free HEK-293, B16F and A375 cells were maintained in DMEM supplemented with 10% fetal bovine serum and penicillin/streptomycin. To measure phosphorylation of IGPR-1 in response to EGF stimulation, cells were plated in 60-cm plates with 10% FBS DMEM overnight with approximate 80–90% confluency. The cells were washed twice with PBS and stimulated with EGF as described in the figure legends. The cells were lysed, and the whole cell lysates were mixed with sample buffer (5×) and boiled for 5 min. The whole cell lysates were subjected to western blotting and immunoblotted with antibody of the interest as described in the figure legends. In some experiments, the cells were treated with a specific chemical inhibitor or transfected with a particular construct as indicated in the figure legends.

### Purification of GST-fusion proteins and GST pulldown assay

Recombinant GST-fusion proteins were purified from the BL21(DE3) Escherichia coli transformed with pGEX-4T2 constructs. Single colony was grown in 5 mL Luria-Bertani (LB) medium overnight at 37°C. The culture was then expanded into −200-300 mL LB medium until an optical density of 0.6. The protein expression was induced by 0.1 mM isopropyl-β-D-thiogalactoside (IPTG) at 30°C for overnight. The cells were collected and re-suspended in GST buffer (25 mM Tris pH 8.0, 5 mM dithiothreitol (DTT), 150 mM NaCl) and sonicated. The GST-fusion proteins were purified via glutathione Sepharose beads. GST pulldown assay was performed as previously described^16^.

### In vitro kinase assay

The Src kinase and EGFR kinase assays were performed as described previously^16^. Briefly, EGFR or c-Src kinase were incubated with the GST-IGPR-1 with 50μL reaction buffer (20 mM PIPES, pH 7.0, 10 mM MnCl2, and 0.2 mM ATP). After 30 min of incubation at 30°C, reactions were terminated by the addition of 2× SDS sample buffer, and samples were subjected to SDS-PAGE followed by western blotting using anti-phospho-Y222 antibody.

### Immunopaired Antibody Detection Analysis

The ActivSignal assay examines phosphorylation or expression of 70 different signaling proteins, which covers 20 major signaling pathways as previously described^17–18^.

### 3D-spheroid cell culture

For 3D-spheroid culture, cells were mixed in approximately 1% methylcellulose and cultured in ultra-low attachment (ULA) 24-well plates (Corning, cat #3473) supplemented with 10%FBS for a duration of 4-8 days or as described in the figure legends.

### 3D modeling of IGPR-1

The alpha-fold predicated structure of the human IGPR-1 was downloaded from the publicly available PDB site. The visualization of the IGPR-1 structure was carried out via PyMOL software (version 4.6.0).

### Melanoma tissue array and immunofluorescence staining

Paraffin-embedded melanoma tissue arrays with 27 cases of melanoma along with adjacent skin and benign nevus (cat# SK181 and T382c) were purchased (Biomax, USA). Except for four patients, all twenty three subjects were in stage IIB/IIC and consisted of 19 females and 8 males. Tissue arrays after deparaffinization and antigen retrieval were permeabilized with 0.3% Triton X-100 in PBS and blocked with 5% BSA. Slides were incubated in primary antibodies (anti-IGPR-1 antibody, 1:200) and phospho-Y222-IGPR-1 antibody, 1:100) overnight at 4°C. After washing, slides were incubated with Alexa Fluor Plus 594 goat anti-rabbit (Invitrogen) antibody for 45 minutes at room temperature. Slides were mounted using ProLong Gold Antifade and DAPI and imaged using Nikon Deconvolution Wide-Field Epifluorescence. ImageJ (NIH version 2.14.0/1.54f) was used to quantify IGPR1 and pY222-IGPR-1 levels in melanoma. Image J quantifies the signal using integrated density, which is a combination of pixel intensity and pixel number and has been used in the past to quantify expression of various proteins in tissues^19–20^. Twenty-seven individual melanoma array were examined. ROIs were drawn in three high-power field images per section and average quantification was made.

### Statistical analysis

Statistical analysis was performed using GraphPad Prism 8 (GraphPad Software, La Jolla, CA) or SAS version 9.3 SAS Institute, Cary, NC). Results are expressed as the mean ± SEM or median, range, and 25^th^ and 75^th^ percentile in box and whisker plots depending on the normality of data. Student’s t-test was performed for group comparison. For multiple groups, overall group comparison was first examined using analysis of variance (ANOVA). With the rejection of null hypothesis, the post-hoc analysis using Bonferroni correction or the pairwise comparisons with Tukey’s multiple comparison procedure were performed. P values of ≤ 0.05 were considered statistically significant.

## Results

### Identification of ITAM motif in the cytoplasmic domain of IGPR-1

The immunoreceptor tyrosine-based activation motif (ITAM) and its variant form is called immunoreceptor tyrosine-based inhibitory motif (ITIM), is a loosely conserved sequence of amino acids that appears in the sequence of Y-x-x-I/L (where x is any amino acids) and is found in the cytoplasmic tails of diverse group of receptors, including immune cell receptors, cell adhesion molecules and others^21–24^. ITIM consensus sequence is originally described as S/I/V/L-x-Y-x-x-I/V/L (where x is any amino acids)^25^, Phosphorylation of the conserved tyrosine residue in the ITAM/ITIM motif recruits the SH2 domain-containing proteins, leading either to activation or inhibition of the receptor function^21, 26–27^. A closer look at the sequence homology of ITAM/ITIM motif in the CD28 family proteins indicates that the ITAM/ITIM motif represents a divergent sequence surrounding a key tyrosine phosphorylation residue (**Figure 1A**).The cytoplasmic tail of IGPR-1 contains two putative tyrosine phosphorylation sites within the ITAM/ITIM motifs, VLY_197_RPR and SIY_222_STS (**Figure 1A**). The predicted structure of IGPR-1 shows that the key residues in these motifs are orientated differently (**Figure 1B, C**), which could influence their recognition by the tyrosine kinases and the SH2 domain-containing substrates. Furthermore, the VLY_197_RPR motif is located in the structured region, whereas the SIY_222_STS motif is not, which could make the VLY_197_RPR motif potentially less accessible to the substrates and the upstream tyrosine kinases.

**Figure 1:**
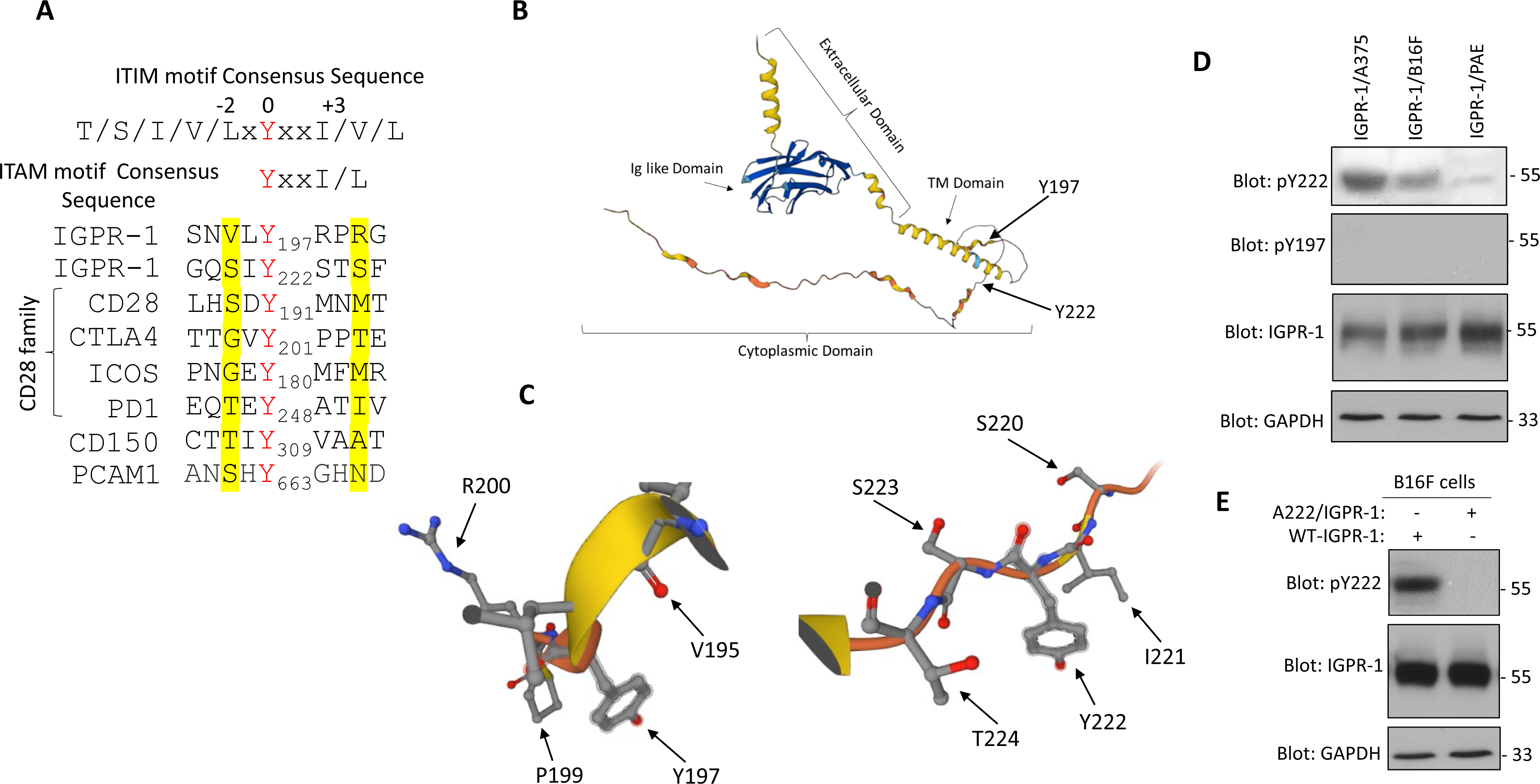
immunoreceptor tyrosine-based motif phosphorylation of IGPR-1. (**A**) An ITAM/ITIM-like motif is present in the cytoplasmic domain of IGPR-1. Amino acid sequences of ITAM/ITIM in CD28 family receptors and unrelated receptors. (**B, C**) Predicated structure of IGPR-1 and the locations of Y197 and Y222. The alpha-fold predicated structure of the human IGPR-1 was downloaded from the PDB site. The visualization of the IGPR-1 was carried out via PyMOL software. (**D**) Whole cell lysates from A375, B16F, and PAE cells ectopically expressing IGPR-1 were subjected to western blot analysis. (**E**) Mutation of Y222 abolishes the detection of IGPR-1 by phospho-222 antibody.

To investigate the phosphorylation of IGPR-1 at Y197 and Y222, we developed rabbit polyclonal anti-phospho-Y197 and anti-phospho-Y222 antibodies. These antibodies specifically recognized the corresponding phosphorylated Y197 and Y222 peptides in a in ELISA assay (data not shown). Next, we analyzed phosphorylation of Y197 and Y222 via phospho-Y197 and phospho-Y222 antibodies in multiple cell lines ectopically expressing IGPR-1. IGPR-1 was strongly phosphorylated at Y222 only in melanoma cell lines, including the human skin melanoma cell line (A375) and mouse skin melanoma (B16F) cell line, but very weakly in PAE cells (**Figure 1D**). Phosphorylation of Y197 in the same cell lysates was not detected (**Figure 1D**), indicating that Y197 is not phosphorylated, or its detection is beyond the sensitivity of this antibody. To further test the cell type-dependent phosphorylation of Y222, we examined phosphorylation of Y222 in colon cancer cell lines, HT29 and HCT116 ectopically expressing IGPR-1. However, we did not detect phosphorylation of Y222 in these cell lines, (**S. Figure 1**), indicating that Y222 is preferably phosphorylated in the melanoma cells but not in colon cancer cell lines or in endothelial cells. Mutation of Y222 abolished the recognition of IGPR-1 by phospho-Y222 antibody (**Figure 1E**), indicating that the anti-phospho-Y222 antibody specifically recognizes phosphorylated Y222. Taken together, we show that IGPR-1 is phosphorylated at Y222 in a context-dependent manner.

### IGPR-1 is upregulated and phosphorylated at Y222 in human melanoma

Next, we asked whether IGPR-1 is phosphorylated at Y222 in human melanoma. We examined expression and phosphorylation of Y222 of IGPR-1 in twenty seven melanoma cases via immunofluorescent staining. Except for four patients, all the twenty three subjects were in stage IIB/IIC. Adjacent normal skin and nevi were used as a control. Negative controls for both the antibodies showed no staining (**S. Figure 2**). The staining of melanoma tissues with the previously validated IGPR-1 antibody^2, 5^ showed that IGPR-1 is upregulated in melanoma compared to control adjacent tissue (**Figure 2A**). More importantly, IGPR-1 was also strongly phosphorylated at Y222 (**Figure 2B**). Phosphorylation of Y222 in fifteen melanoma cases (55%) were higher than the median range (**Figure 2B**). Of these melanomas, eight melanomas showed stronger positive signal, which was higher than the 75^th^ percentile. Interestingly, although IGPR-1 expression pattern in melanoma was variable (**Figure 2C**), however, overall, its expression was increased from 2.8 to 4-fold in melanoma compared to adjacent skin (P=0.0287) and nevus (P =0.0054). Nearly 60% (16 out of 27) melanomas showed higher IGPR-1 levels which was above the 75^th^ percentile of adjacent skin and nevus. The results showed that in human melanoma, IGPR-1 is both upregulated and phosphorylated at Y222.

**Figure 2.**
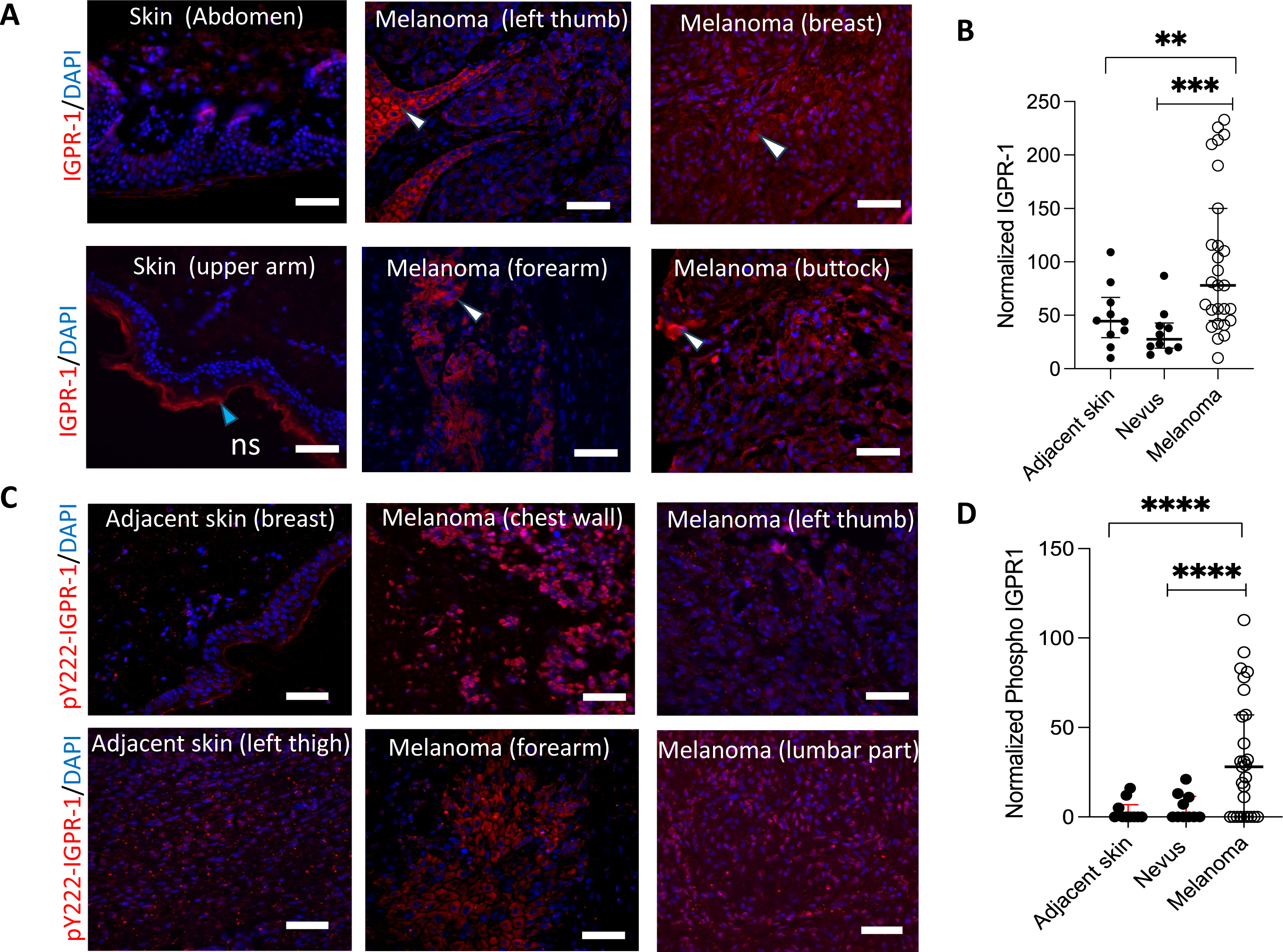
IGPR-1 is expressed and phosphorylated at Y222 in melanoma. (**A**). Representative images of expression of IGPR-1 in human skin melanoma, adjacent skin and benign nevi are shown. The white arrowhead points to the IGPR-1 positive cells in melanoma. Scale bar 100 microns. (**B**). The normalized integrated density of IGPR-1. The lines correspond to the median, 25^th^, and 75^th^ percentile. Ordinary one-way ANOVA was used to compare all the groups (P = 0.0029). Student’s t test was performed to compare two groups. (**C**). Representative images of IGPR-1 phosphorylation at Y222 in melanoma, adjacent skin and benign nevi are shown. DAPI stains nuclei. Scale bar 100 microns. (**D**). The normalized integrated density of pY222. The lines correspond to the median, 25^th^, and 75^th^ percentile. Ordinary one-way ANOVA was used to compare all the groups (P = 0.0027). Student’s t-test was performed to compare two groups.

Given the observed results, we analyzed human skin melanoma genome sequencing dataset (Broad/Dana Farber study, 2012)^28^ via online cBioport^29–30^ (accessed on Feb 10, 2024). Our analysis revealed that IGPR-1 is mutated in 7.69% of melanoma cases (n=26) (**S. Figure 3A**). Analysis of several other genome sequencing datasets of human skin melanomas also uncovered various alterations (ranging from 5.26%-1.2%) in the coding region of IGPR-1, including mutations, deep deletion, and amplification (**S. Figure, 3A**). Summary of IGPR-1 mutations identified in human skin melanomas is shown (**S. Figure 3B**). Mutation of G218 (shown in the red box, **S. Figure 3B**) could affect phosphorylation of IGPR-1 at Y222 but this was not investigated in this study. Our further analysis of the TCGA dataset showed that IGPR-1 mRNA is upregulated in 5% (n= 419 cases) of primary skin melanoma (**S. Figure 3C**). We asked whether IGPR-1 mutations correlate with the overall survival of patients with melanoma. Our Kaplan-Meier survival analysis on the same datasets revealed that the overall survival of patients with IGPR-1 mutations was lower (median survival 49.27 months vs. 61.55 months) compared to the patients without IGPR-1 mutations (**S. Figure 3D**). Taken together, IGPR-1 is mutated and upregulated in melanoma, and is distinctively phosphorylated at Y222.

**Figure 3:**
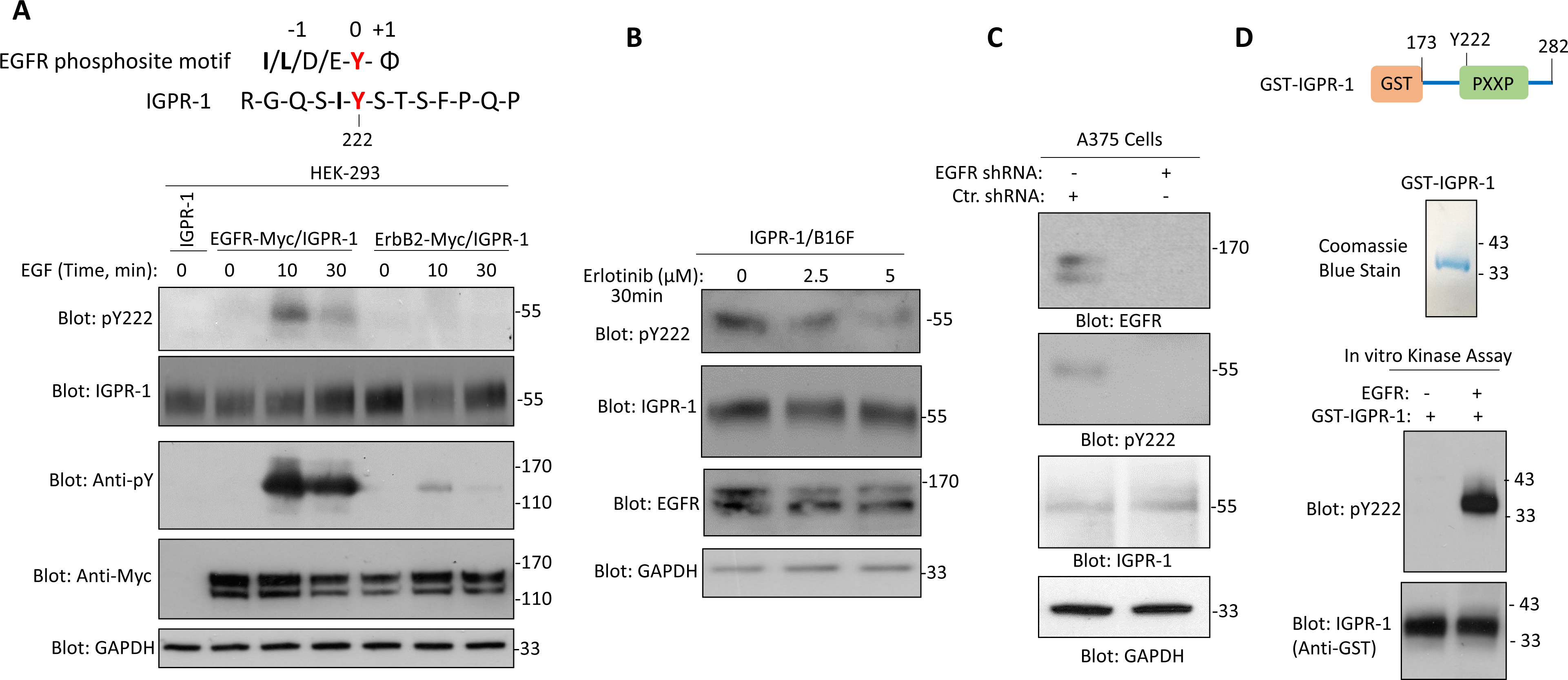
EGFR phosphorylates IGPR-1 at Y222. (**A**) The conserved amino acid sequence of EGFR phosphorylation motif. φ, hydrophobic amino acids. (**B**) HEK-293 cells stably expressing IGPR-1 were transfected with EGFR or ErB1. After 48 hours, cells were stimulated with control vehicle or EGF for the indicated times and cells were lysed and whole cell lysates were subjected wot western blot analysis. EGFR but not ErbB2 stimulated phosphorylation of IGPR-1 at Y222. (**C**) B16F cells expressing IGPR-1 were treated with the EGFR inhibitor (erlotinib) with various concentrations and cells were lysed after 30 minutes and whole cell lysates were analyzed for phosphorylation of IGPR-1 via western blot analysis. (**D**) A375 cells were treated with human EGFR shRNA and after 48 hours cells were analyzed for phosphorylation of IGPR-1 at Y222. (**E**) Schematic of GST-fusion IGPR-1 and Coomassie blue stain of the purified GST-IGPR-1. In vitro kinase assay is performed as described in the material and methods section. The result shows that recombinant EGFR phosphorylates GST-IGPR-1 at Y222.

### EGFR and Src kinase phosphorylate IGPR-1 at Tyrosine 222

EGFR is known to preferentially phosphorylate its substrates at the tyrosine site with Ile/Leu or acidic residues (*e.g.* Glu, Asp) before the tyrosine (−1 position) and a hydrophobic residue (*e.g*. Ala, Ile, Leu, Met, Phe, Val, Pro or Gly) in the +1 position^31–32^. EGFR strongly favors an Ile/Leu residue at the −1 position^31^. The −1 position of Y222 of IGPR-1 also is Ile (I) (**Figure 3A**), suggesting that EGFR could phosphorylate Y222. To this end, we co-expressed EGFR or ErbB2/EGFR2 with IGPR-1 in HEK-293 cells and monitored the phosphorylation of IGPR-1 at Y222 via western blot analysis. We chose HEK-293 cells because Y222 is not phosphorylated in HEK-293 cells and its phosphorylation is only induced in response to EGFR stimulation. As shown, stimulation of cells co-expressing EGFR with IGPR-1 with EGF resulted in the phosphorylation of Y222 but not in the cells co-expressing IGPR-1 with ErbB2 (**Figure 3A**). Furthermore, phosphorylation of Y222 in B16F cells was inhibited with the EGFR-specific inhibitor erlotinib^33^ in a dose-dependent manner (**Figure 3B**). Erlotinib is in a clinical trial for the treatment of cutaneous squamous cell carcinoma^34^. Next, we showed that knockdown of EGFR in human A375 melanoma cells inhibits the phosphorylation of endogenously expressed IGPR-1 (**Figure 3C**). To demonstrate the direct role of EGFR in the phosphorylation of Y222, we generated recombinant GST fusion IGPR-1 encompassed of cytoplasmic domain (**Figure 3D**) and demonstrated that a highly purified recombinant active EGFR catalyzed phosphorylation of GST-IGPR-1 at Y222 (**Figure 3D**), supporting the direct role of EGFR in the phosphorylation of Y222. Given that EGFR and the Src family kinases often select similar substrates for phosphorylation ^31, 35^ and Src family kinases are also known to phosphorylate the tyrosine in the ITIM/ITAM consensus sequence motif ^36–37^, we also tested whether c-Src can phosphorylate Y222. Under identical *in vitro* kinase assay conditions, both EGFR and c-Src kinase phosphorylated Y222, albeit c-Src catalyzed phosphorylation of Y222, was significantly less than that of EGFR (**S. Figure 4**), indicating that both EGFR and c-Src kinase could phosphorylate IGPR-1 at Y222.

**Figure 4.**
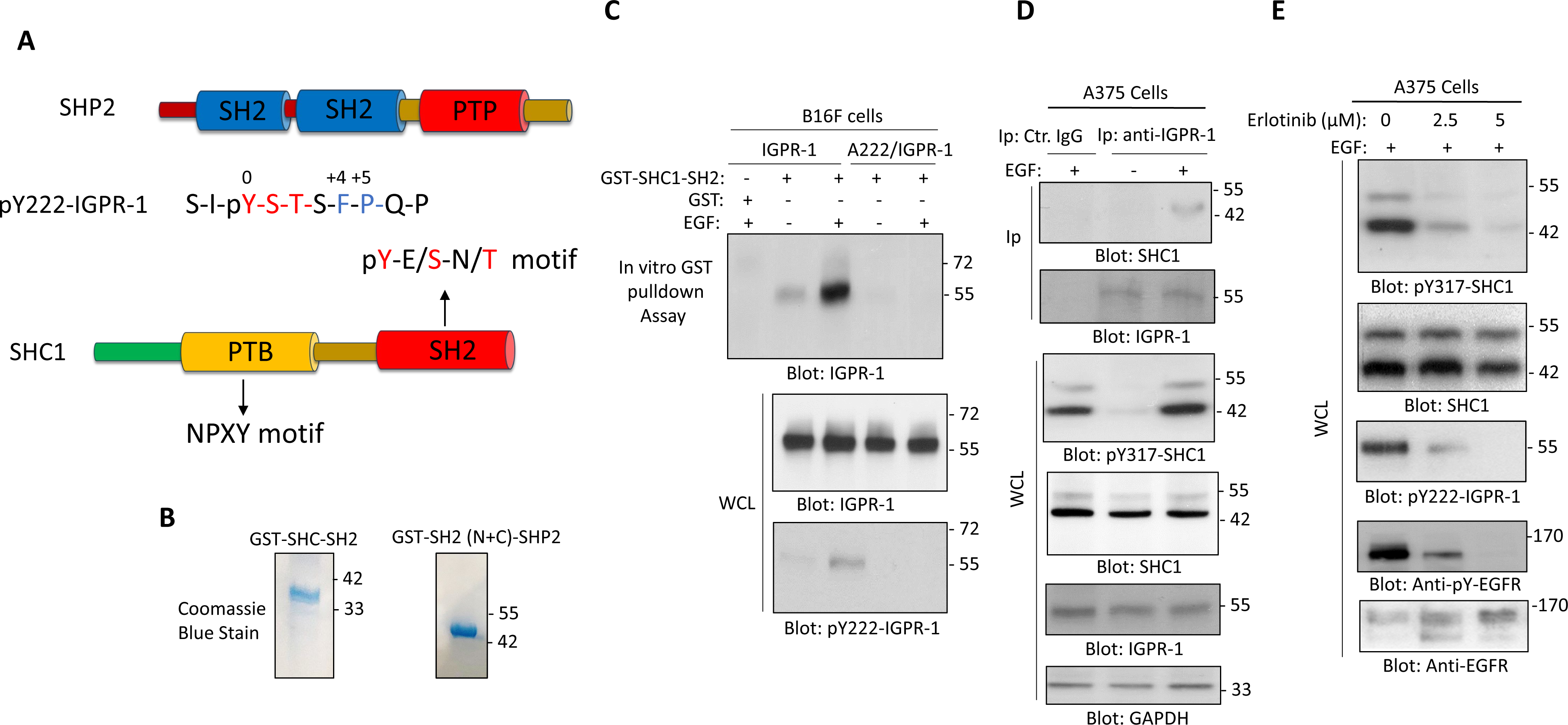
Phosphorylation of Y222 mediates the interaction of SHC and SHP2 with IGPR-1. (**A**) Schematics of SHC1 and SHP2 and their preferred binding consensus sequences. (**B**) The Coomassie blue staining of the purified GST-SH2-SHC and GST-SH2 (N+C)-SHP2. (**C**) B16F cells expressing IGPR-1 or A222-IGPR-1 were stimulated with the control or EGF and whole cell lysates were subjected to GST-pulldown assay using GST-SHC as a bait, as described in the material and methods section. (**D**) Whole cell lysate from A375 endogenously expressing IGPR-1 was subjected co-immunoprecipitation assay via anti-IGPR-1 antibody. IGPR-1 interacts with GST-SH2-SHC stimulated with EGF. (**E**) A375 cells were treated with EGFR inhibitor (erlotinib) and phosphorylation of SHC and IGPR-1 were determined via western blot analysis.

### Phosphorylation of Y222 mediates the binding of IGPR-1 with the SH2 domain containing proteins SHC1 and SHP2

Phosphorylation of the tyrosine in the ITIM/ITAM motifs of various receptors commonly known to recruit the SH2 domain-containing proteins such as SH2 domain-containing phosphatase 2 (SHP2) and others, leading to a diverse array of signal transduction mechanisms^38–39^. The SH2 domain of SHP2 is shown to prefer hydrophobic aromatic residues (*e.g.,* Trp, Tyr, and Phe) at the +4, and +5 positions^40–41^. The amino acids at the +4 and +5 positions of phospho-Y222 are also hydrophobic aromatic residues (**Figure 4A**). Additionally, the phospho-Y222 also could interact with the SH2 domain of Src homology 2 domain-containing-transforming protein C1 (SHC1). The SH2 domain of SHC1 preferentially recognizes the phospho-tyrosine sites encompassing pY-E/S-N/T motif ^42^. As shown, the phospho-Y222 shares a similar consensus sequence motif (**Figure 4A**). First, we examined whether the SH2 domain of SHC1 interacts with IGPR-1 via an *in vitro* GST pulldown assay using GST-SH2-SHC1 (**Figure 4B**). The result showed that the GST-SH2-SHC1 interacted with IGPR-1 and its binding with IGPR-1 further increased with EGF stimulation (**Figure 4B**). Mutation of Y222 to Alanine (A222) totally abolished the interaction of IGPR-1 with the GST-SH2-SHC1 (**Figure 4B**), underscoring the importance of phospho-Y222 in the recognition of the GST-SH2-SHC1 protein.

Furthermore, we also carried out an *in vivo* binding assay via co-immunoprecipitation using whole cell lysate of A375 cells endogenously expressing IGPR-1 and SHC1 and showed that IGPR-1 co-immunoprecipitated with SHC1 in an EGF-stimulated manner (**Figure 4D**). EGF stimulation of A375 cells also promoted phosphorylation of SHC1 (**Figure 4D**). Furthermore, blocking EGFR via erlotinib in A375 cells inhibited both SHC1 and IGPR-1 phosphorylation in a dose-dependent manner (**Figure 4E**), further supporting the central role of EGFR in the activation of IGPR-1. Additionally, we asked whether the extracellular domain of IGPR-1 which mediates its homophilic dimerization^5^, affect its interaction with SHC1. GST-pulldown assay showed that deletion of the homophilic dimerization domain abolishes the binding of IGPR-1 with the SH2-SHC1 (**S. Figure 5**). This deletion also inhibited phosphorylation of IGPR-1 at Y222 (**S. Figure 5**), indicating that the homophilic dimerization of IGPR-1 is a prerequisite for the EGFR to catalyze the phosphorylation of IGPR-1 at Y222.

**Figure 5.**
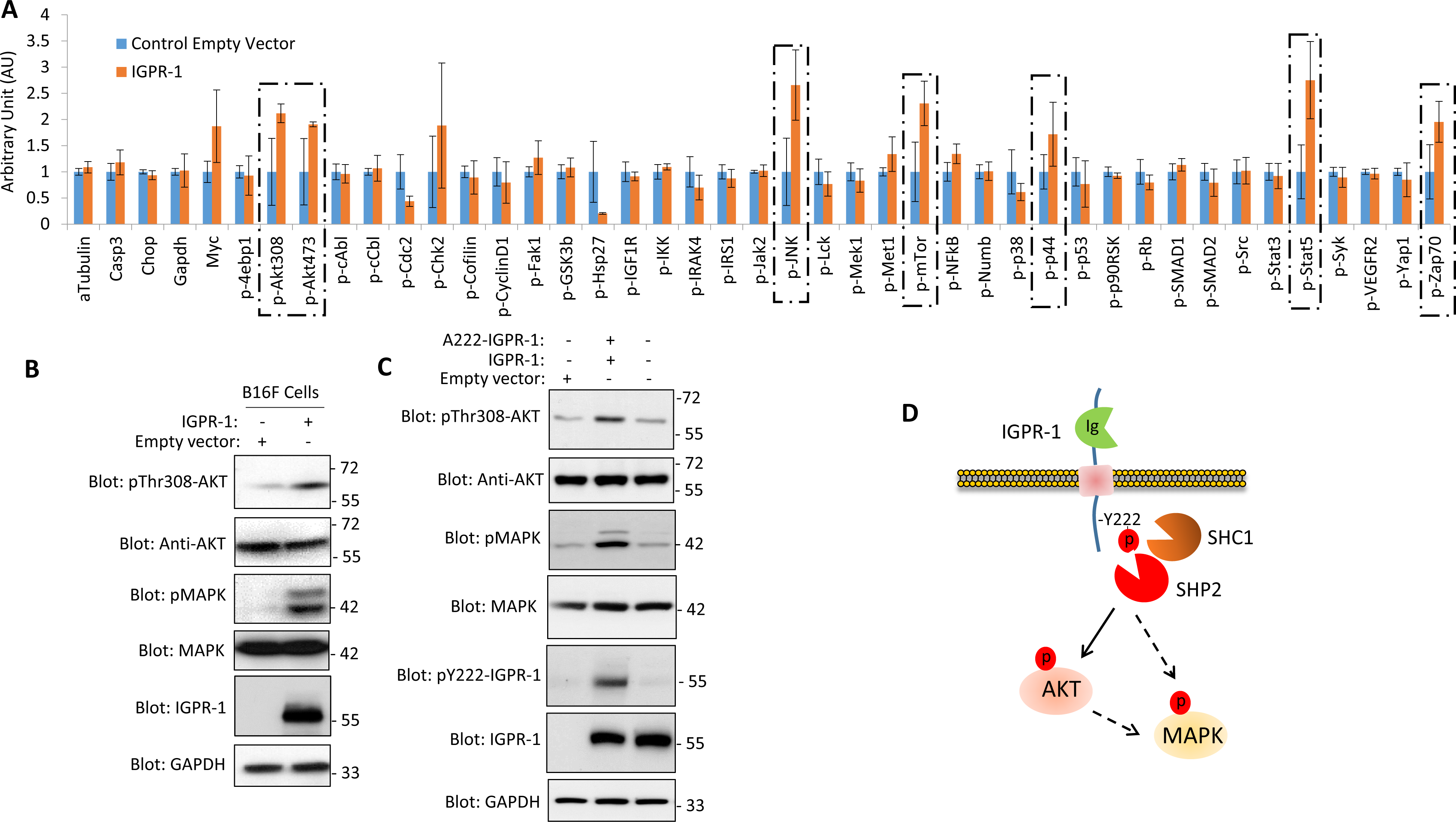
IGPR-1 activates multiple signaling pathways. (**A**) B16F cells expressing empty control vector or IGPR-1 were plated in 96-well plates in triplicates and subjected to ActiveSignal Assay analysis, as described in the material and methods section. The graph shows the activation of major pathways affected by IGPR-1 (dotted boxes). AU (arbitrary unit). (**B**) Whole cell lysate from B16F cells expressing control vector or IGPR-1 was subjected to western blot analysis using a specific antibody, as indicated. (**C**) whole cell lysates from B16F cells expressing IGPR-1 or A222-IGPR-1 were subjected to western blot analysis and phosphorylation of AKT and MAPK were determined. (**D**) Schematic of pY222-dependent activation of AKT and MAPK.

### IGPR-1 stimulates activation of AKT pathway and supports 3D-spheroid tumor growth

To elucidate the signaling mechanism of IGPR-1, we analyzed the activation of twenty major pathways consisting of 70 individual signaling proteins (the effect of IGPR-1 in the activation of forty individual signaling proteins shown) via immunopaired antibody detection system (ActiveSignal Assay) analysis platform^17–18^. Among the major pathways that were stimulated by IGPR-1 in B16F cells were AKT, mTOR, MAPK (ERK1/ERK2), JNK, STAT5, and ZAP70 (**Figure 5A**). Activation of these signaling pathways in various cell types and cancers is shown to stimulate cell proliferation and survival^43–46^. IGPR-1-dependent activation of AKT is particularly important as AKT is known to activate mTORC1 through phosphorylation of tuberous sclerosis complex 2 (TSC2) and PRAS40^47^ and MAPK by phosphorylation of Raf1^48^. We further validated the activation of AKT and MAPK by western blot analysis. As shown, AKT and MAPK were selectively phosphorylated in IGPR-1/B16F cells but not in the control cells (**Figure 5B**). Next, we investigated if the activation of AKT and MAPK in B16F cells by IGPR-1 is influenced by the phosphorylation of Y222. Western blot analysis revealed a significant decrease in the phosphorylation of AKT and MAPK in A222-IGPR-1/B16F cells compared to B16F cells expressing the wild-type IGPR-1 (**Figure 5B**), indicating that activation of AKT and MAPK by IGPR-1 is dependent on the phosphorylation of Y222.

In our previous report, we demonstrated that IGPR-1 accelerates tumor growth by facilitating the formation of multicellular aggregation in colon cancer cells^9^. Hence, we performed tests to assess the potential of IGPR-1 in promoting tumor growth within a 3D-spheroid culture. The result showed that B16F cells expressing empty vector (EV) uniformly formed 3D spheroids relatively similar size. However, B16F cells expressing IGPR-1 developed varying sizes ranging from medium, large, and very large spheroids, but overall, they were larger compared to EV/B16F spheroids (**Figure 6A**). The spheroid formation of B16F cells expressing the A222-IGPR-1 was significantly reduced compared to IGPR-1/B16F cells (**Figure 6A**), showing that Y222 phosphorylation plays a critical role in the tumor spheroid growth of B16F cells. We further examined whether IGPR-1-mediated spheroid growth is linked to its ability to promote cell survival. Results from propidium iodide (PI) staining of B16F cells revealed that EV/B16F spheroids exhibited high cell death (PI positive) after 24 hours, while IGPR-1/B16F spheroids showed minimal cell death (**Figure 6B**). The result also showed that A222/IGPR-1 spheroids were largely dead (**Figure 6B**), indicating that phosphorylation of Y222 plays an important role in supporting B16F cell survival in spheroid conditions. To investigate the effect of AKT inhibition on 3D-spheroid growth, we treated IGPR-1/B16F cells with a selective AKT inhibitor (MK-2206). This was done considering the fact that IGPR-1 activates AKT and the mutation of Y222 on IGPR-1 blocks AKT activation. MK-2206 treatment significantly inhibited spheroid growth (**Figure 6C**), indicating the important role of AKT in IGPR-1-induced spheroid growth.

**Figure 6.**
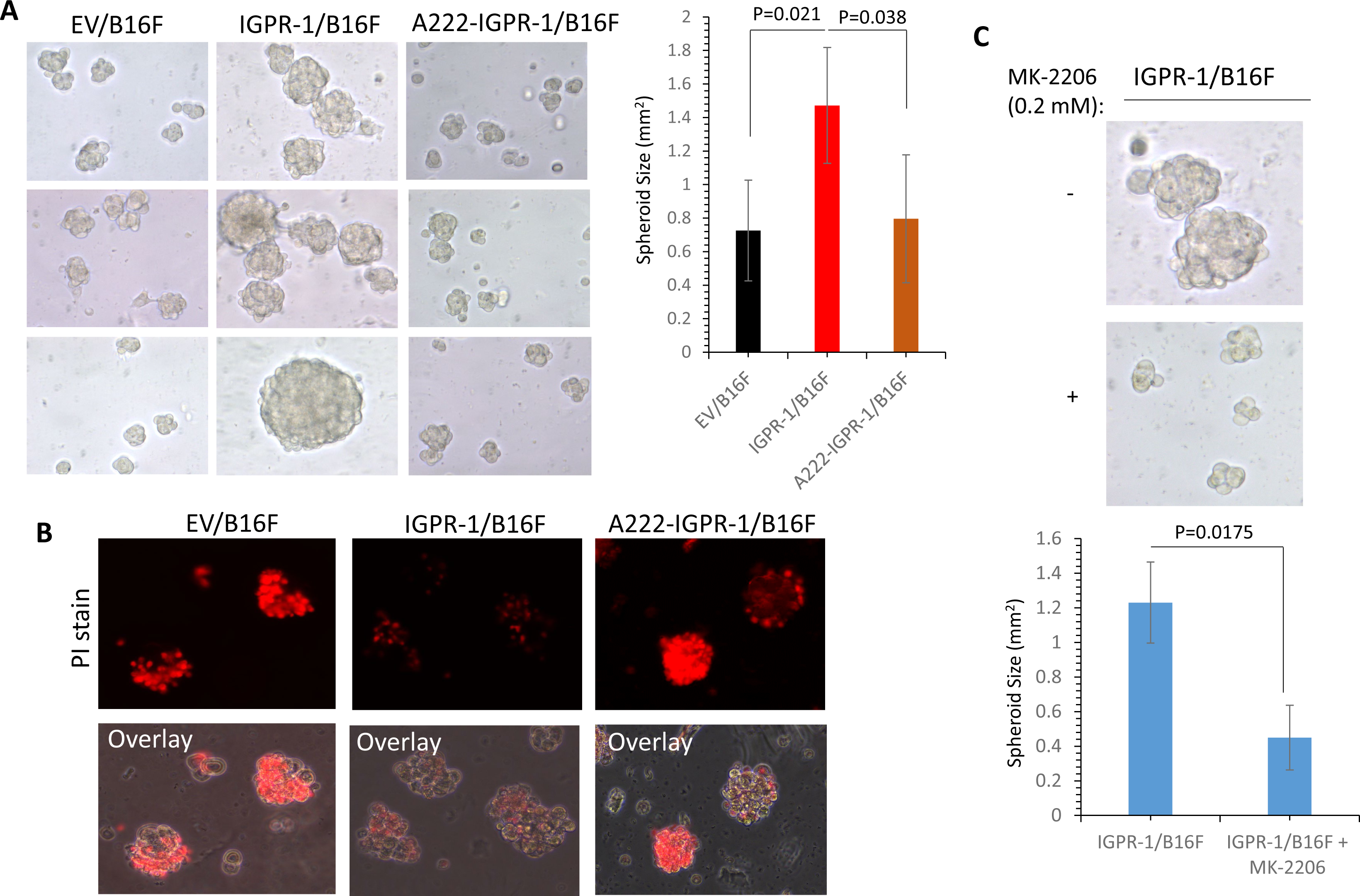
Phosphorylation of Y222 supports 3D-spheroid growth of B16F melanoma cells. (**A**) Spheroid growth of B16F cells expressing empty vector (EV), IGPR-1, or A222-IGPR-1 after four days. The graph represents at least twenty spheroids per group of triplicates. Very large spheroids from the IGPR-1/B16F group was excluded from quantification because it was giving a large standard deviation. (**B**) Staining of 3D-spherioids with propidium Iodide. (**C**). Spheroids of IGPR-1/B16F cells were treated with AKT inhibitor (MK-2206) or control vehicle. The graph represents twenty spheroids per group of triplicates.

## Discussion

The data presented in this study provide evidence for the phosphorylation of IGPR-1 at Y222 in melanoma and its role in the recruitment of SHP2 and SHC, which leads to the activation of multiple signaling pathways including AKT in melanoma cells. Recruitment of SHC and SHP2 with various receptors has been shown to stimulate MAPK and AKT pathways^49–52^. ITAM/ITIM-based tyrosine phosphorylation and signaling are versatile in their ability to select a particular SH2 domain. In some cases, such as CD28 (Y_191_MNM), CTLA4 (Y_201_VKM), and ICOS (Y_180_MFM) the ITIM/ITAM-based tyrosine phosphorylation results in the interaction with the SH2 domain of p85/PI3 kinase^53^, which is consistent with the known conserved preference of the SH2 domain of p85 for YxxM motif (where x is any amino acid)^54^. In some other cases, such as PD1 phosphorylation of ITAM/ITIM tyrosine recruits SHP2 to the receptor^55^. More recent study demonstrated that the ITIM/ITAMs interact with a broad array of SH2 domains^38^. Our results demonstrated that the ITAM phosphorylation of IGPR-1 recognizes the SH2 domains of SHC1 and SHP2. However, the mode of the recognition of phosphorylated Y222 by the SH2 domains of SHC1 and SHP2 could be different. The SH2 domain of SHC1 is known to select hydrophobic amino acids ((E/D/N/S/T) at +1 and +2 positions^42^. The amino acids at +1 and +2 positions of pY222 also are hydrophobic amino acids. On the other hand, the SH2 domains of SHP2 prefer a large hydrophobic residue (W,Y, F) at +4 or +5 positions^40^. The amino acids at +4 and +5 positions after pY222 are F and P (**Figure 1A**). *Ptpn11* encodes SHP2 and it acts as a positive regulator of cell proliferation and anti-apoptosis by stimulating the Ras/MAPK pathway^56^. SHC1 is an SH2 domain adapter protein that is also known to interact with various cell surface receptors, including RTKs such as EGFR, which leads to cell proliferation via activation of AKT and MAPK pathway^57^.

We found both EGFR and Src phosphorylate IGPR-1 at Y222. EGFR is hyperactivated in various cancer types, including melanoma ^58–61^ and its hyperactivation in melanoma is associated with tumor progression and poor survival ^60, 62–64^. EGFR and Src phosphorylate substrates that are highly enriched in sequences that conform to the −1 acidic and +1 hydrophobic residues^31^. Src kinase also is activated by EGFR and other RTKs and non-RTK receptors^65^ and mediates tyrosine phosphorylation of ITAM/ITIM in other receptors^37, 66^.

Our study showed an upregulation of both mRNA and protein levels for IGPR-1. Surprisingly, the IGPR-1 protein exhibited a more pronounced upregulation than its mRNA. Although the mRNA levels usually do not correspond to protein levels, the dysregulation of mechanisms governing IGPR-1 in melanoma at the protein level could account for these differences. We have previously reported that NEDD4 binds to IGPR-1 and promotes ubiquitination-dependent degradation of IGPR-1^11^. Downregulation of NEDD4 expression or its activity in certain melanoma cases could lead to increased expression of IGPR-1. IGPR-1 also is mutated in 7.69% of melanoma cases. However, it is currently unclear how these mutations affect IGPR-1 function in melanoma. Further studies are required to fully elucidate the complexity of IGPR-1 mutation/upregulation and the role of Y222 phosphorylation-dependent signaling in melanoma.

Our analysis on the signaling pathways initiated by IGPR-1 point toward the unique pro-survival and anti-apoptosis function of IGPR-1 in melanoma cells. IGPR-1 expressed in mouse B16F melanoma cells stimulated AKT, mTOR, JNK, STAT5, and ZAP70 pathways, which are all key pro-tumor pathways. Activation of AKT, mTOR, JNK, and MAPK are strongly linked to various aspects of tumorigenesis including, cellular proliferation and metabolic programing of cancer cells^67–69^. Mutation of Y222 abrogated activation of AKT and MAPK, indicating that phosphorylation of Y222, in part, plays a central role in orchestrating the pro-tumor activity of IGPR-1 in melanoma cells. In support of the pro-survival function of IGPR-1 in melanoma, IGPR-1 stimulated 3D-spheriod growth of B16F cells and blocking AKT activity or mutation of Y222 on IGPR-1 abolished its pro-tumorigenic activity.

### Conclusions

The work presented here demonstrated the upregulation and phosphorylation of IGPR-1 in melanoma. IGPR-1 is selectively phosphorylated at Y222, and upon phosphorylation by EGFR and Src kinase, it recruits SHP2 and SHC, leading to the activation of AKT and MAPK pathways. Phosphorylation of Y222 on IGPR-1 and activation of AKT are required for IGPR-1-dependent 3D-spheroid tumor growth.

### Limitation of the study

IGPR-1 stimulates various key pro-tumor signaling molecules, including AKT, mTOR, and MAPK pathways in melanoma. However, further studies are required to link mechanistically to the phosphorylation of IGPR-1 at Y222. Moreover, the role of IGPR-1 in melanoma needs further investigation. However, because IGPR-1 is not present in mice^2^, in vivo studies of IGPR-1 present a serious challenge. IGPR-1 is mutated or upregulated in human skin melanoma, which could lead to its hyperactivation. However, further studies are required to establish a link between IGPR-1 mutation and/or over-expression in the pathobiology of melanoma.

## Declarations

### Ethics Approval and Consent Participation

No human or animals involved in this study.

### Consent for publication

The author agrees with the published version of the manuscript.

### Availability of data and materials

Cell lines, plasmids and other reagents described in this manuscript are available at a reasonable request.

### Competing interests

The author declares no competing interest.

### Funding

This work was partially supported by the Center of Cross Organ Vascular Pathology, Department of Medicine, Boston University Medical Center.

### Author’s Contributions

NR is responsible for conceptualization, methodology, formal analysis, data curation and writing the manuscript. VC examined the human tissue and analyzed the data, edited the ms. SP stained the melanoma cores.

## Acknowledgements

none

## Notes

**Conflict of Interest**: The author declares no conflict of interest.

### Competing Interest Statement

The authors have declared no competing interest.

