## Supplemental data for "IGPR-1 is phosphorylated on the immunoreceptor tyrosine-based motif, stimulates AKT pathway and supports melanoma growth"

**S. Figure 1. IGPR-1 is not phosphorylated at Y222 in endothelial and colon cancer cell lines**. Whole cell lysates from PAE cell (porcine aortic endothelial cells), colon cancer cell lines (HCT116 and HT29) ectopically expressing IGPR-1 or Serine 220 mutant IGPR-1 (A220) were lysed and subjected to western blot analysis.

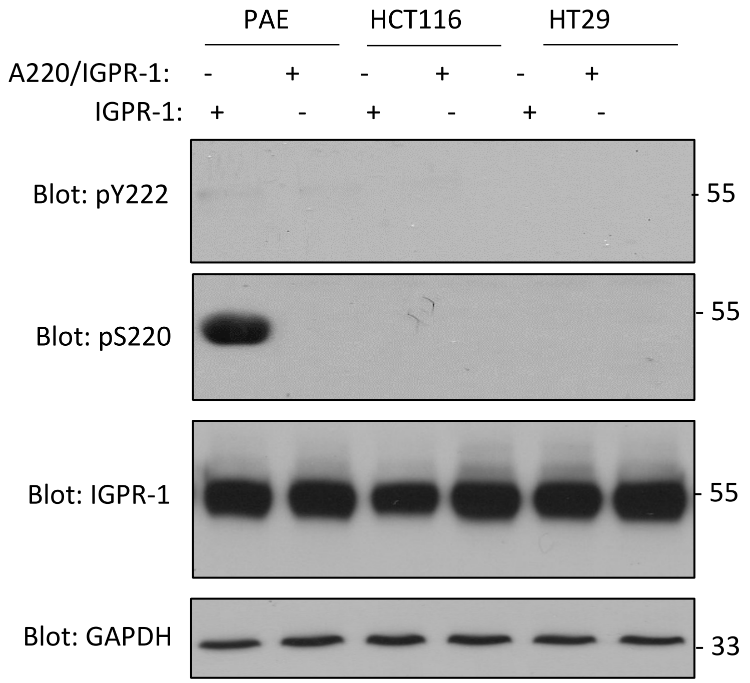

**S. Figure 2. Negative staining for IGPR-1 and PY222-IGRPR-1**

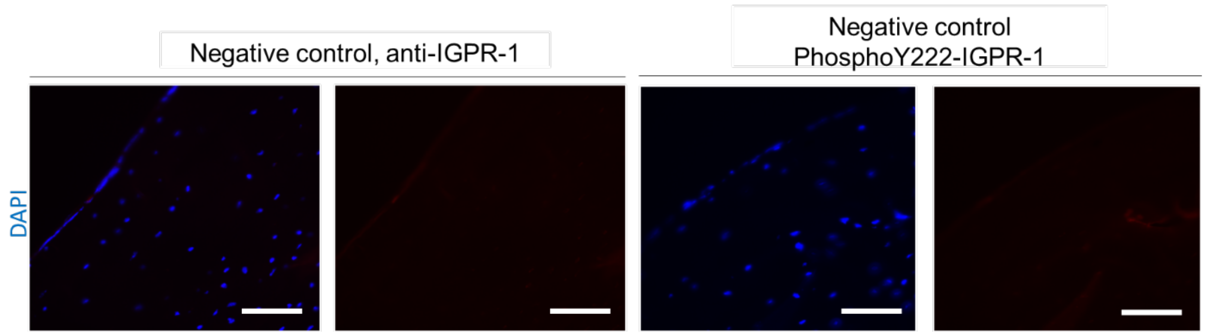


**S. Figure 3. Src kinase phosphorylates IGPR-1 at Y222**. Equal amount of EGFR and Src kinase recombinant proteins were added to an in vitro kinase assay and phosphorylation of IGPR-1 at Y222 was determined in western blot analysis using anti-pY222 antibody. The graph is a representative of a triplicate experiments.

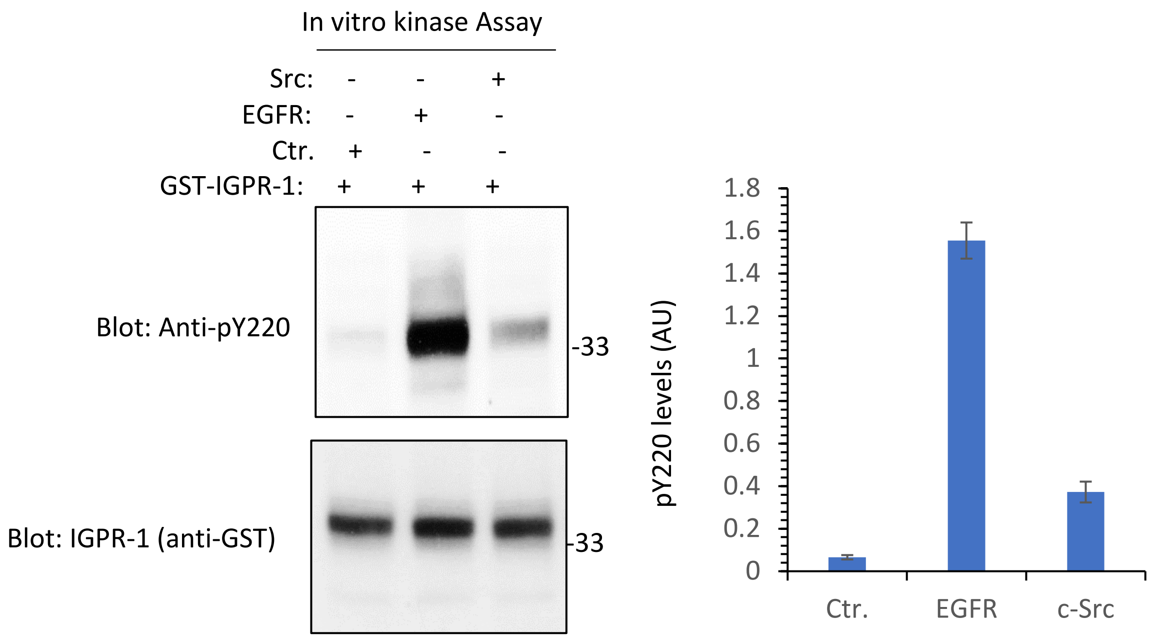

**S. Figure 4. The extracellular domain of IGPR-1 is required for its phosphorylation at Y222**. Whole cell lysates from B16F cells expressing control empty vector, IGPR-1, or the extracellular domain truncated IGPR-1 (ΔN-IGPR-1) were analyzed for phosphorylation of IGPR-1 at Y222 via western blot analysis. WCL, whole cell lysate.

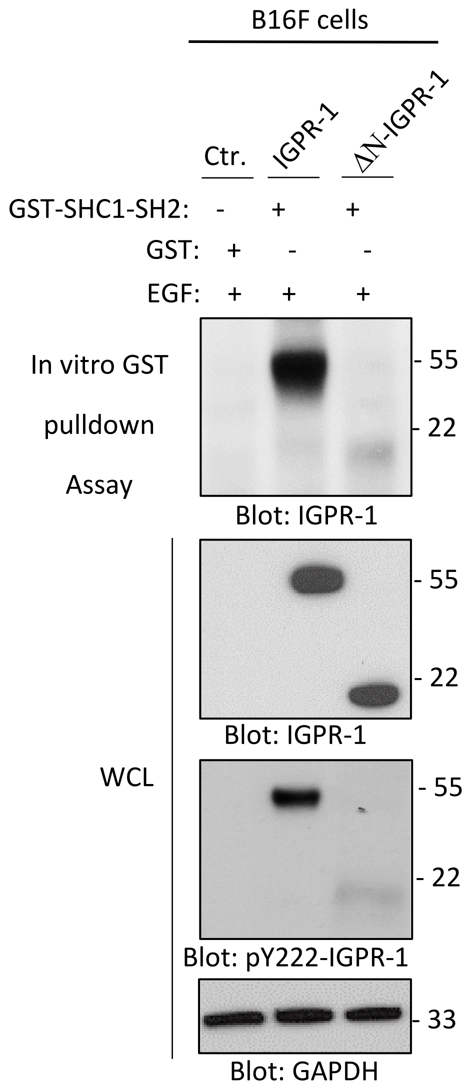

**S. Figure 5. IGPR-1 is frequently mutated and upregulated in human skin melanoma and its mutation correlates with the overall poor survival.** (**A**) Frequency of IGPR-1 mutations in human skin melanomas (various studies). (**B**) Summary of IGPR-1 mutations in human skin melanoma. (**C**) Volcano plot showing IGPR-1 mRNA is upregulated in 5% (n=419 cases) of primary human skin melanoma (data extracted from the TCGA dataset, 02/12/2024). (**D**) Kaplan-Meier survival analysis.

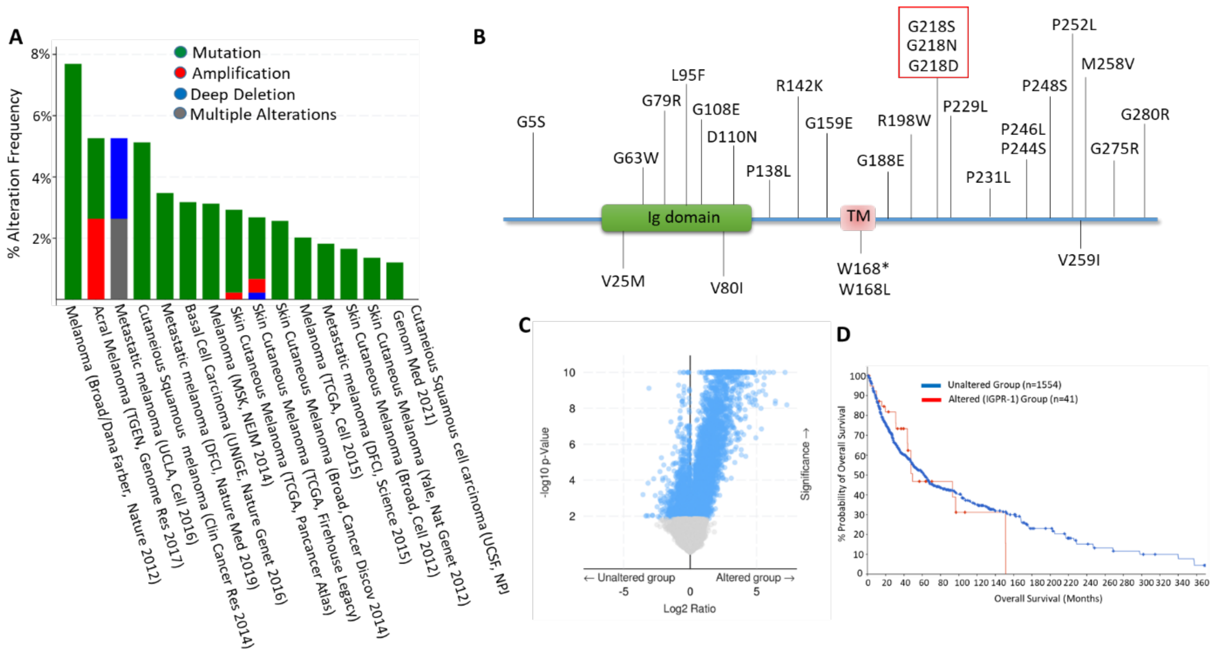
